# An EMG Foundation Model for Neural Decoding

**DOI:** 10.64898/2025.12.17.694831

**Authors:** Andrew Garrett Kurbis, Alex Mihailidis, Brokoslaw Laschowski

## Abstract

Decoding algorithms can be used to predict motor behaviour from patterns of neural activity. However, most studies rely on subject-optimized models, limiting generalization and scalability to novel subjects and tasks. Building on recent advances in deep learning and large-scale data, here we developed an EMG foundation model for neural decoding. Our model was trained on more than 197 hours of neural recordings from 1,667 subjects. We used unsupervised learning to pretrain our encoder layers on unlabeled data, followed by supervised learning on our benchmark dataset. Additionally, we performed large-scale architecture searches to develop a custom encoder-decoder model composed of convolutional and transformer layers, optimized for both scalability and performance. Our foundation model consistently outperformed the previous state-of-the-art (i.e., subject-optimized models) across both in-distribution and out-of-distribution evaluations. For in-distribution evaluation, few-shot fine-tuning yielded an average F1 score of 0.697, compared to 0.638 for subject-optimized models. For out-of-distribution evaluation on clinical and demographically-shifted subjects, we achieved an average F1 score of 0.599, compared to 0.518 for the subject-optimized baselines. Taken together, our results highlight the value of foundation models for robust and generalizable neural decoding. By publicly releasing our neural network weights and training pipeline, we aim to support future research in computational neuroscience and neural-machine interfaces.

## I. Introduction

**N**EURAL decoding is the process of predicting behaviour from patterns of neural activity [1], which can be recorded, for example, using surface electromyography (EMG). However, variability between subjects, sensor placement, and physiological factors can lead to significant differences in neural signals [2]. As a result, most state-of-the-art models are optimized for individual subjects and/or tasks. Many applications in neural-machine interfaces and computational neuroscience require decoding algorithms with high accuracy and generalization [3], [4]. Learning at scale (i.e., using large datasets, models, and compute) has the potential to transform the field of neural decoding, akin to what foundation models have done in other domains.

This challenge is often described as *domain shift*. Here, we define a domain as a feature space *X* and joint probability distribution *P* (*X, Y*), where *X* are the input features and *Y* are the behavioural labels. Domains differ when their feature spaces or joint distributions differ, which can hinder generalization, especially out-of-distribution.

Although advances in machine learning have improved neural decoding, most models are optimized for individual subjects and/or tasks. Both achieve strong in-distribution accuracy but generalize poorly to new subjects, sessions, or environments. For example, [5] achieved 96% decoding accuracy with an adaptive Bayesian model, but required retraining for each subject. Krausz et al. [6] reported 92% accuracy in steady state tasks and 75% accuracy in transition states using linear discriminant analysis (LDA), but generalization remained difficult even within small groups. Task-optimized models, such as [7], have shown high decoding accuracy in controlled environments, but have lacked evaluation under domain shift or at scale.

Foundation models offer a promising solution for generalization in neural decoding [8], [9]. This new class of models uses unsupervised pretraining, large and diverse datasets, and deep neural networks to support effective transfer learning. Foundation models have shown strong generalization in other domains [8]–[13], but their development and use for neural decoding is still emerging.

In this study, we present our development of an EMG foundation model for neural decoding (Figure 1). To develop and train our model, we developed one of the largest EMG datasets published to date, comprising over 1,600 subjects and 200 hours of neural recordings. Our custom encoder-decoder architecture, which combines convolutional and transformer layers, was benchmarked against state-of-the-art models optimized for individual subjects. By publicly releasing our neural network weights and training pipeline, we aim to support future research in computational neuroscience.

**Fig. 1.**
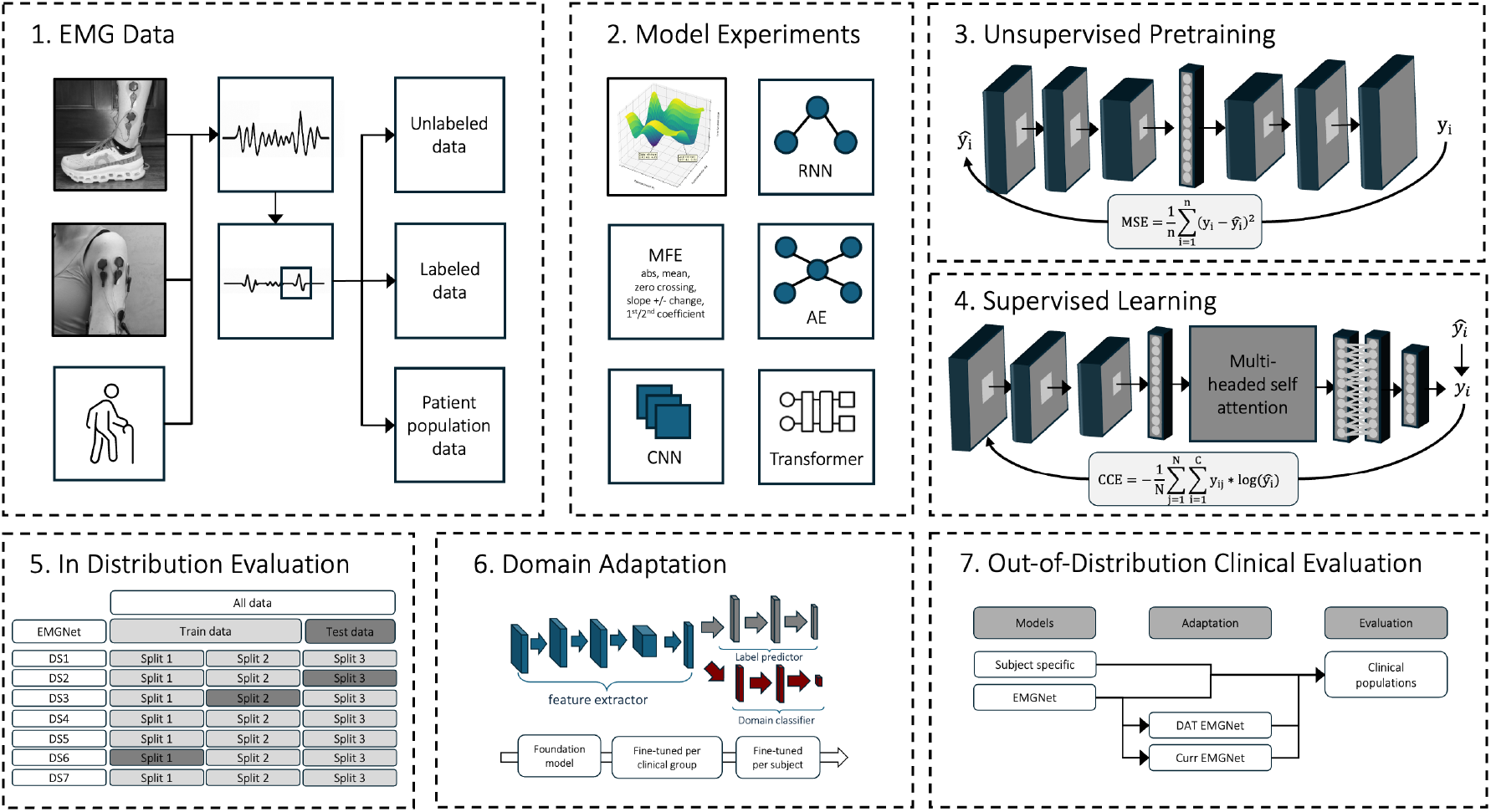
Summary of our foundation model development and evaluation, including (1) development of unlabeled, labeled, and patient population datasets; (2) evaluation of our neural network architecture and feature processing; (3) unsupervised pretraining; (4) supervised learning; (5) in-distribution evaluation using 3-fold cross-validation; (6) domain adaptation at group level; and (7) out-of-distribution evaluation on clinical and demographically-shifted subjects.

## II. Methods

### A. Dataset

To develop and test our foundation model, we developed one of the largest EMG datasets published to date, comprising 11,809 minutes (197 hours) of neural recordings from 1,667 subjects, aggregated across 43 open-source datasets. Data were organized into three functional subsets for (1) unsupervised pretraining, (2) supervised learning on our benchmark decoding tasks, and (3) out-of-distribution evaluation on clinical and demographically-shifted subjects (see Figure 2).

**Fig. 2.**
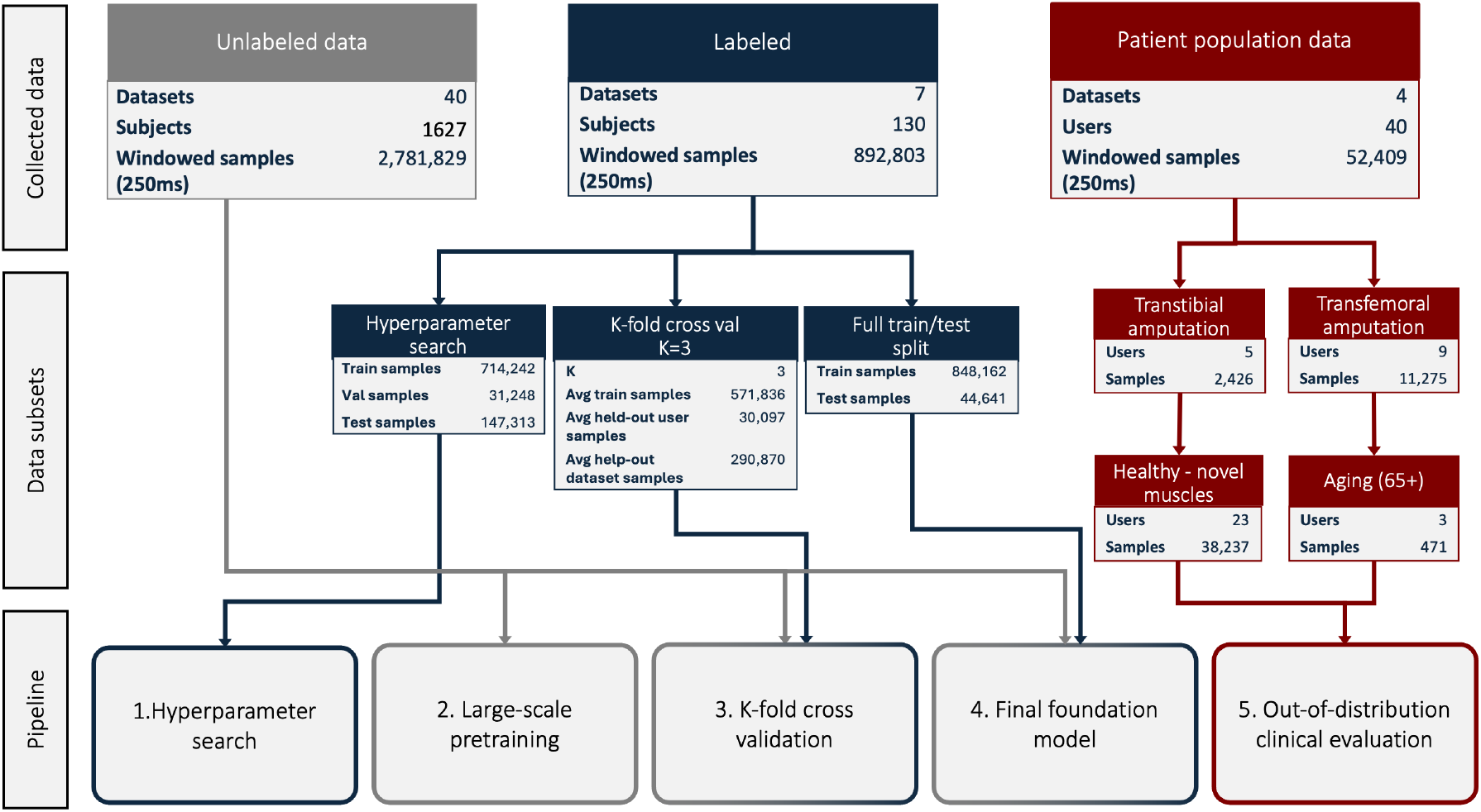
Data used to develop and evaluate our foundation model, including (1) large unlabeled dataset for unsupervised pretraining, (2) labeled dataset for model selection, supervised learning, and in-distribution evaluation, and (3) patient population dataset for testing out-of-distribution.

#### 1) Unlabeled data

To support generalization, we developed a large unlabeled dataset for unsupervised pretraining. This subset includes 11,500 minutes of unlabeled data from 1,627 subjects across 40 datasets, 18 muscles, and 10 tasks. The absence of behavioural labels or experimental constraints enabled our neural network to learn generalizable representations from a wide range of subjects and tasks, which subsequently served as inductive biases in our optimization.

#### 2) Labeled data

For supervised learning, we developed a labeled dataset containing 3,500 minutes of data and 890,000 labeled windows from 132 subjects [14]. We focused on the tibialis anterior, medial gastrocnemius, rectus femoris, and biceps femoris, given their relevance to our proof-of-concept decoding tasks and consistent availability across datasets. A wrapper-based ablation confirmed these muscles were the most informative such that removing them caused the largest drops in accuracy. The label space included six different motor tasks: standing, walking, stair ascent and descent, and ramp ascent and descent. Some subjects were included in both labeled and unlabeled datasets.

#### 3) Patient population data

For testing out-of-distribution, we developed a dataset with 218 minutes of neural recordings from 40 subjects across three axes of domain shift: (1) clinical variation (patients with amputation [5]), (2) sensor variation (healthy subjects with novel muscles [15]), and (3) demographic variation (older adults, 65+ [16]). These variants introduced fundamentally different data distributions, which enabled principled testing of our model’s robustness to domain shifts. To evaluate generalization, we compared the subject-optimized models trained from scratch against our pretrained foundation model with few-shot fine-tuning.

#### 4) Signal processing

We developed a signal processing pipeline in accordance with best practices [17]. Data were resampled to 1000 Hz, bandpass filtered (20–500 Hz), notch filtered (50/60 Hz), rectified, and normalized using mean dynamic method. Data were segmented into 250 ms windows with 30 ms stride and labeled using the final frame. Fixed dataset splits (80%/3.5%/16.5%) were used for pretraining and supervised learning, with leave-one-subject-out cross-validation for testing generalization.

### B. Foundation Model

We performed a large hyperparameter search using our labeled dataset to identify the optimal neural network architecture. Our search evaluated different combinations of encoder and decoder models, with joint optimization of training hyperparameters to ensure robust performance. This process enabled selection of an optimal architecture for both supervised and unsupervised learning, supporting generalization across diverse subjects and tasks.

We formulated our benchmark decoding tasks as a supervised sequence classification problem. Given an input sequence **x** ∈ ℝ^*T* ×*C*^, where *T* is the number of time steps and *C* the number of channels, the goal was to learn a model *f*_*θ*_ parameterized by *θ* that predicts the behavioural label *y ∈ *y**. During training, the model minimizes the empirical loss over the labeled dataset 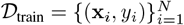:

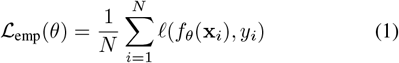

where *ℓ* (·, ·) is the categorical cross-entropy loss. We included *ℓ*_2_ regularization to prevent overfitting. Although the labeled dataset was fixed, it contains samples from multiple subjects *u*= { *u*_1_, …, *u*_*M*_}, each with distinct data distributions. The goal of our architecture search was to identify a neural network that minimizes the expected loss across this set of subject-specific sub-distributions. Here, *D* _*u*_ is the subset of training samples from subject *u*. The optimization objective is:

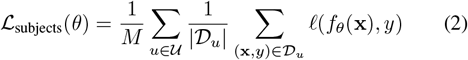

This encouraged the neural network to capture transferable features across subjects, enabling the model to generalize to novel subjects without retraining. As such, we did not optimize our neural network for a single subject or task-specific distribution, but rather for robust generalization across subject variability within the labeled dataset.

We explored three encoding methods in our architecture search: (1) manual feature extraction (time-domain autoregressive features: mean absolute value, zero crossings, waveform length, slope sign changes, and autoregressive coefficients), (2) direct time-series input (raw data), and (3) convolutional neural networks (temporal CNNs with ReLU and batch normalization). We also studied three decoding methods: (1) feedforward neural networks, (2) LSTM recurrent neural networks, and (3) transformer attention-based models. All encoders and decoders were designed to be scalable in depth, width, and number of layers.

To compare the encoder and decoder models, we performed an optimization sweep over the training and regularization hyperparameters, including learning rate, dropout rate, batch size, normalization, and oversampling. Our goal was to isolate the architecture performance from confounding effects due to suboptimal training dynamics (see Table I).

**TABLE I.** The search space of training and architecture hyperparameters in our system optimization.

| Parameter | Search space |
| --- | --- |
| Window Size | 250 ms raw EMG window (250x4) |
| Latent Dim | Embedding size $\in \{6, 16, 32, 64\}$ |
| Conv Filters | 32–256 filters; optionally doubled across layers |
| Kernel Size | $\{3, 5, 7, 15, 25, 50, 75\}$ |
| FNN Layers | 1–16 layers; 32–512 neurons each |
| LSTM Units | 1–8 layers; 8–128 hidden units |
| Transformer Config | 1–8 layers; 32–256 dim; 1–8 heads; FFN: 64–512 |
| Loss Function | Sparse CE, Weighted CE, Focal Loss |
| Oversampling | Random oversampling with noise and jitter |
| Batch Size | $\{128, 256, 512, 1024\}$ |
| Learning Rate | $1 \times 10^{-5}$ to $5 \times 10^{-3}$ (log scale) |
| Dropout | Rates from 0.1 to 0.5 |
| L2 Weight Decay | $\{0, 10^{-4}, 10^{-3}, 10^{-2}\}$ |
| Optimizer | Adam, AdamW |

We evaluated each model using three metrics: (1) F1 score, which was our main metric chosen for its ability to balance precision and recall and robustness to class imbalance; (2) accuracy, which we included for consistency with previous studies; and (3) number of learnable parameters, which serves as a proxy for model size and efficiency.

We developed a four-stage pipeline to test each hyperparameter combination (see Figure 3). We ran preliminary runs to define feasible hyperparameter bounds, followed by Bayesian optimization sweeps using Weights and Biases. Our top-performing models were retrained with early stopping to ensure convergence and generalization. All training was performed on a multi-GPU cluster and runtime and memory efficiency were monitored for real-time and embedded applications.

**Fig. 3.**
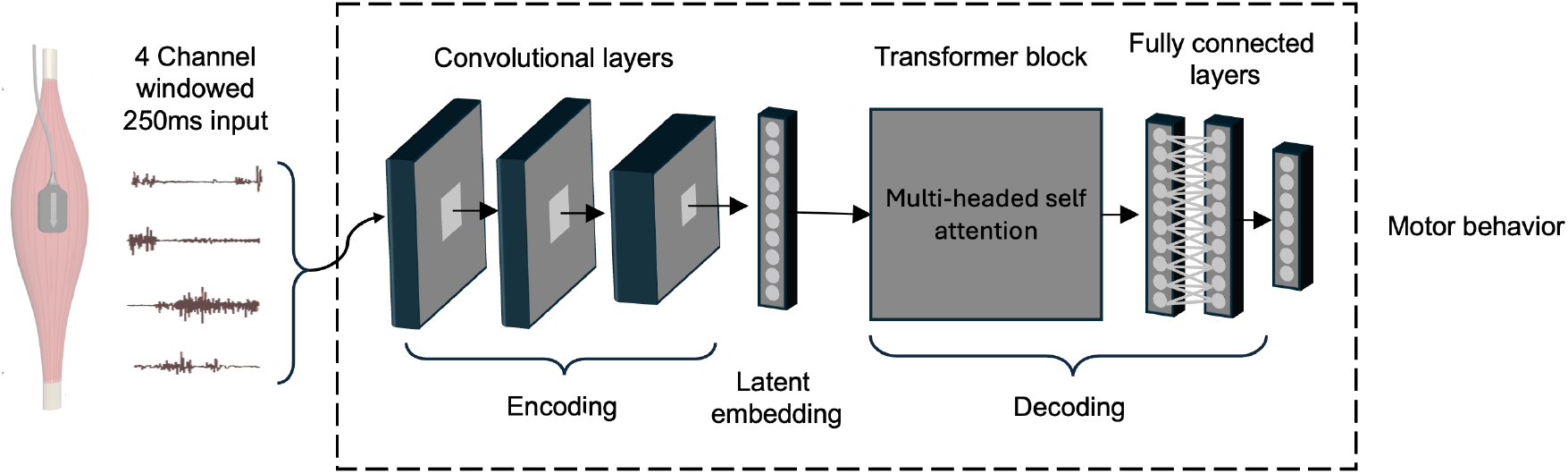
Our foundation model architecture, including (1) convolutional encoder for local feature extraction, (2) transformer decoder for modeling long-range dependencies, and (3) two fully connected layers for classification/decoding. This architectural design balances efficiency, computational feasibility, and representational capacity.

Our search identified the optimal architecture to be a convolutional encoder, transformer decoder, and two fully connected layers. Neural networks using raw data, especially CNN encoders, significantly outperformed traditional handcrafted features, and transformers achieved the best performance among all decoders. The CNN-transformer combined model was the highest performing overall, achieving statistically significant improvements over baseline manual feature extraction. This compact yet expressive architecture (1.75 million parameters) balances efficiency, computational feasibility, and representational capacity. All code, sweeps, and model checkpoints are available at https://github.com/gkurbis/EMGNet for reproducibility.

### C. Subject-Optimized Models

We used subject-optimized models as our baseline for comparison. These models were optimized using subject-specific labeled data without pretraining, shared representations, or access to population-level information. We used the same neural network architecture as previously described, but performed hyperparameter optimization for each individual subject.

This hyperparameter search was also used to determine the training configuration for fine-tuning our foundation model. By using the subject-optimized models for the baseline and fine-tuning, we biased the comparisons in favor of these models, ensuring that any performance gains observed from our foundation model could not be attributed to biased hyperparameters.

To develop the subject-optimized models, we explored the following search space: learning rate (1×10^−4^ to 1×10^−2^),batch size (128 to 512), dropout rate (0.1 to 0.5), and L2 weight decay (1 × 10^−5^ to 1 × 10^−3^). Each model was trained from scratch and evaluated using the same performance metrics as previously described. These subject-optimized models, representative of the current state-of-the-art, provided an empirical upper bound for neural decoding performance without transfer learning and pretrained weights.

### D. Model Training

#### 1) Unsupervised learning

We used the unlabeled dataset to optimize our encoder model to learn robust and transferable representations. To decouple representation learning from task-optimized objectives, we adopted a reconstruction pretraining strategy. This enabled our model to learn intrinsic patterns in the neural activity without relying on behavioural labels, forming a generalizable embedding space for downstream decoding tasks. The unsupervised pretraining objective was to minimize the mean squared error between the original and reconstructed signals, where *x*_*n,t,c*_ and 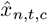 are the original and reconstructed neural signals, and *N, T, C* are batch size, time steps, and channels.

#### 2) Supervised learning

Using supervised learning, we then adapted the pretrained encoder, transformer decoder, and classification layers to our benchmark decoding tasks using the labeled dataset. The supervised learning objective was to minimize the class-weighted categorical cross-entropy loss:

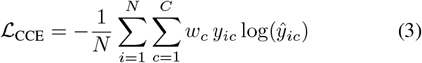

where *w*_*c*_ is the class weight, *y*_*ic*_ is the one-hot target, and 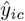 is the predicted probability. To study representational transfer, we varied the number of encoder layers that were frozen. Freezing only the first convolutional layer provided the optimal balance such that early layers preserved general representations learned during pretraining, while deeper layers remained free to adapt to task-specific representations.

#### 3) K-fold cross validation

We used 3-fold cross-validation to test in-distribution. This enabled robust benchmarking of our foundation model on left-out subjects and datasets. Both zero-shot and few-shot generalization were evaluated, with and without the pretrained encoder, to quantify the impact of the unsupervised pretraining. Finally, we trained our foundation model on the full labeled dataset with a 10% validation split and early stopping, using the optimal hyperparameters from the cross-validation. Our neural network weights and scripts are available at https://github.com/gkurbis/EMGNet for reproducibility.

### E. Domain Adaptation

To support out-of-distribution generalization, we studied two population-level adaptation techniques: (1) domain adversarial training and (2) curriculum learning. Our goal was to reduce distributional shifts between the training and target populations while preserving decoding performance.

#### 1) Domain adversarial training

We used domain adversarial training to learn domain-invariant neural representations across source domains (healthy subjects) and target domains (clinical and demographically-shifted subjects). Similar to [18], we appended a domain discriminator to our pretrained encoder and added a gradient reversal layer in-between. This gradient reversal layer multiplies the incoming gradient by −*λ*, forcing the encoder to confuse the discriminator and align distributions across domains. The total loss minimized is:

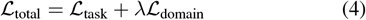

where ℒ_task_ is the weighted categorical cross-entropy loss for neural decoding, ℒ_domain_ is the binary cross-entropy loss for domain classification (source vs. target subjects), and *λ* is a scalar controlling the trade-off between decoding accuracy and domain alignment. If *G*_*f*_ is the feature encoder, *G*_*y*_ the label predictor, and *G*_*d*_ the domain discriminator, then the training minimizes the empirical risk:

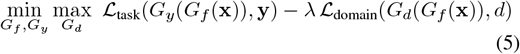

where **x** are the EMG windows, **y** are the behavioural labels, and *d* ∈ { 0, 1} is the domain label (0 = source, 1 = target). The encoder model *G*_*f*_ is co-optimized to perform accurate neural decoding and to fool the discriminator *G*_*d*_ into predicting the domain labels incorrectly.

We performed separate domain adversarial training for each shifted subgroup (e.g., amputees and older adults), treating each as an independent target domain. We co-optimized the encoder and discriminator using mixed-domain mini-batches, and used the adapted encoder for subject-level fine-tuning. We then evaluated domain alignment using target-domain classification accuracy and AUC metrics on the discriminator.

#### 2) Curriculum learning

We also studied curriculum learning, where we fine-tuned our pretrained foundation model to align with the target distribution. In the first stage, we adapted the model by minimizing the task loss:

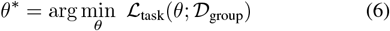

where*D* _group_ is the labeled data from the target domain, excluding the evaluation subject. This tuned the model to domain-specific data statistics. In the second stage, we fine-tuned the model to the evaluation subject using a small subject-specific dataset *D*_subject_ (*M* ≪ *N*):

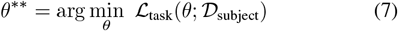

starting from *θ*^*^. This two-stage curriculum learning simulates real-world applications where subject-specific data is scarce but cohort-level data is accessible.

### F. Out-of-Distribution Evaluation

We tested out-of-distribution using clinical and demographically-shifted subsets. Although the output label space remained unchanged, these subsets introduced significant domain shifts. Here, we define out-of-distribution evaluation as the model’s ability to generalize from source domains*D* _*S*_ (healthy subjects) to target domains *D*_*T*_ (clinical or demographically-shifted subjects) where *D*_*S*_ = *D*_*T*_ in data distribution. The marginal input distributions *P*_*S*_(*x*) and *P*_*T*_ (*x*) differ, though the label space *y* remains consistent. Our objective is:

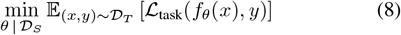

where *f*_*θ*_ is our foundation model pretrained on *D*_*S*_, and ℒ_task_ is the weighted categorical cross-entropy loss. We evaluated four model variants for each out-of-distribution subject: (1) Subject-Optimized: where we trained the model from scratch using the subject’s labeled data; (2) Foundation Model: where we fine-tuned the foundation model on the subject’s labeled data; (3) DAT-Foundation Model: where we adapted the foundation model using domain adversarial training on clinical data, then fine-tuned on the subject; and (4) Curriculum-Foundation Model: where we adapted the foundation model using curriculum learning from the cohort, followed by subject-level fine-tuning.

We evaluated performance using F1 score on the 30% held-out test set from each subject’s *s* labeled dataset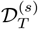. The domain adversarial training and curriculum learning variants used additional group-level data,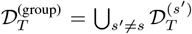, for intermediate adaptation prior to fine-tuning on the evaluation subject. All models were trained using the same 70/30 subject-level train/test split and hyperparameters as previously described. For the domain adversarial training and curriculum learning, we performed an optimization sweep over the number of frozen layers in each neural network, where the freeze depth ranged from 0–3 layers for the CNN encoder, 0–1 layers for the transformer decoder, and 0–1 layers for the classifier.

## III. Results

Our foundation model consistently outperformed the previous state-of-the-art (i.e., models optimized for individual subjects) across both in-distribution and out-of-distribution evaluations. Although our zero-shot foundation model achieved limited (6) generalization on held-out in-distribution subjects (F1 score = 0.505 ±0.049) (see Figure 4), fine-tuning with a small amount of data resulted in significant performance gains (few-shot F1 score = 0.697 ± 0.056), surpassing the subject-optimized models (F1 score = 0.638±0.047) with an average increase in decoding performance of 5.9%.

**Fig. 4.**
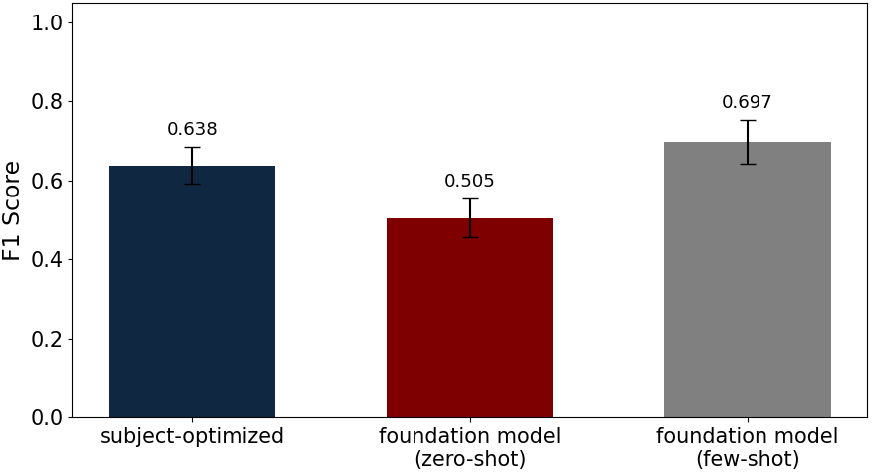
F1 scores for the subject-optimized models, zero-shot foundation model, and few-shot foundation model on held-out in-distribution subjects. Error bars indicate standard deviation across subjects.

With regards to out-of-distribution generalization to domain-shifted subjects (Figure 5), our few-shot foundation model (*F* 1 score = 0.599± 0.102) significantly outperformed the subject-optimized baselines (*F* 1 score = 0.518± 0.180), achieving a 8.1% average increase in decoding performance.

**Fig. 5.**
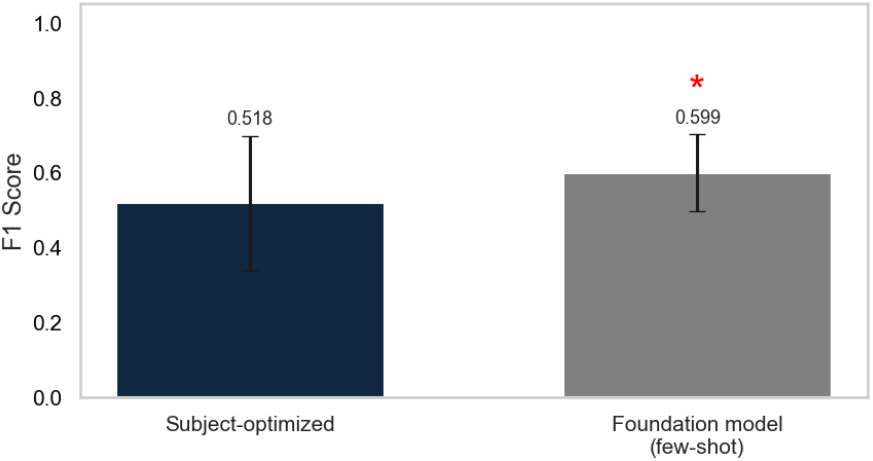
F1 scores for the subject-optimized models and few-shot foundation model on out-of-distribution clinical and demographically-shifted subjects. Error bars indicate standard deviation across subjects.

To quantify the impact of unsupervised learning, we performed an ablation study comparing models with and without pretrained neural network weights. On held-out in-distribution subjects, unsupervised pretraining improved zero-shot generalization from F1 = 0.486 to 0.505. On held-out in-distribution datasets, pretraining improved zero-shot generalization from F1 = 0.345 to 0.387.

Although domain adversarial training achieved an average out-of-distribution F1 score of 0.615±0.116 compared to 0.599±0.102 for our unadapted foundation model (+1.6% improvement), this difference was not statistically significant (*p* = 0.056 with a Wilcoxon signed-rank). Curriculum learning showed no measurable change, achieving an average F1 score of 0.599 in neural decoding.

## IV. Discussion

In this study, we developed an EMG foundation model for neural decoding. To develop and train our model, we created a large-scale dataset with over 1,600 subjects and 200 hours of neural recordings. We used unsupervised learning to pretrain our encoder layers on unlabeled data, followed by supervised learning on our benchmark decoding tasks. We also performed large-scale architecture searches to develop a custom encoder-decoder model composed of convolutional and transformer layers. Our foundation model consistently outperformed the previous state-of-the-art (i.e., subject-optimized models) across both in-distribution and out-of-distribution evaluations, including clinical and demographically-shifted populations. These findings demonstrate the benefit of using pretrained weights compared to subject-optimized models trained from scratch.

Most neural decoding algorithms are optimized for individual subjects and/or tasks (e.g., [5]–[7]). Compared to these subject-optimized models, our foundation model significantly improved performance and generalization. On held-out in-distribution subjects, our few-shot foundation model achieved an average F1 score of 0.697 ± 0.056 compared to 0.638 ± 0.047, *p* < 0.01 for the subject-optimized baselines. This advantage generalized to out-of-distribution populations, achieving an average few-shot F1 score of 0.599 ± 0.102, compared to 0.518 ± 0.180 for the subject-optimized models (*p* = 0.0073). These results highlight the sensitivity of subject-optimized models to domain shift, while our foundation model demonstrated robust generalization.

Compared to task-optimized models [7], our research offers several advantages. Our dataset used for model training comprises more than 200 hours of neural recordings from 1,600 subjects performing various tasks, whereas task-optimized models use far fewer subjects and tasks or are limited to controlled lab environments. Our foundation model learned from natural variability in tasks dynamics, enabling robust representation learning and generalization. This generalization was enabled by unsupervised pretraining–a key differentiator from task-optimized models that rely entirely on supervised learning. By using large unlabeled data from multiple tasks, our foundation model learned more robust representations. Our ablation study showed that using pretrained neural network weights improved zero-shot generalization to novel subjects (from 0.486 to 0.505) and datasets (from 0.345 to 0.387), supporting unsupervised pretraining as a best practice for neural decoding.

Despite these developments, we view our research as a stepping stone toward broader foundation modeling efforts in neuroscience. One promising direction for future research is to study model scaling laws–how changes in neural network size and parameters affect generalization. In other domains, scaling laws have shown predictable improvements with increasing model size until diminishing returns [11], [19]. Although our architecture search included scaling, we did not explicitly evaluate how generalization scaled with the model size. However, we hypothesize that neural decoding will likewise exhibit a capacity threshold.

Lastly, understanding how dataset size and quality affect generalization is critical to advance neural decoding. Previous research in other domains has shown a logarithmic relationship between data quantity and generalization, with diminishing returns as scale increases. Applying these insights to neural decoding could help quantify how much data is needed to improve generalization and whether to prioritize quantity or quality. Furthermore, not all source domains are equally beneficial (i.e., some may introduce negative transfer). Emerging techniques for source selection and subject weighting such as [20] offer promising new strategies for curating training data to optimize performance and generalization.

## V. Conclusion

In conclusion, here we developed an EMG foundation model for neural decoding. By leveraging large-scale, diverse data and unsupervised pretraining, our foundation model consistently outperformed subject-optimized models trained from scratch across both in-distribution and out-of-distribution evaluations– including clinical and demographically-shifted populations– demonstrating improved generalization to novel subjects and tasks. These findings reveal that learned representations from unsupervised pretraining capture shared structure in neural activity that cannot be readily learned from subject-specific data alone. Ultimately, we show that foundation models are a promising new machine learning paradigm for neural decoding, which have both scientific implications for computational neuroscience and practical implications for developing neural-machine interfaces.

## Acknowledgment

This research is dedicated to the students and researchers in Ukraine affected by Russia’s ongoing invasion. Their resilience and unwavering commitment to learning and education serve as a beacon of hope and inspiration for the global academic community.

